# A non-coding A-to-T Kozak site change related to the transmissibility of Alpha, Delta and Omicron VOCs

**DOI:** 10.1101/2021.04.30.442029

**Authors:** Jianing Yang, Guoqing Zhang, Dalang Yu, Ruifang Cao, Xiaoxian Wu, Yunchao Ling, Yi-Hsuan Pan, Chunyan Yi, Xiaoyu Sun, Bing Sun, Yu Zhang, Guo-Ping Zhao, Yixue Li, Haipeng Li

**Author notes:** These authors contributed equally (Jianing Yang, Guoqing Zhang, Dalang Yu, Ruifang Cao).

## Abstract

Three prevalent SARS-CoV-2 Variants of Concern (VOCs) were emerged and caused epidemic waves. It is essential to uncover the key genetic changes that cause the high transmissibility of VOCs. However, different viral mutations are generally tightly linked so traditional population genetic methods may not reliably detect beneficial mutation. In this study, we proposed a new pandemic-scale phylogenomic approach to detect mutations crucial to transmissibility. We analyzed 3,646,973 high-quality SARS-CoV-2 genomic sequences and the epidemiology metadata. Based on the sequential occurrence order of mutations and the instantaneously accelerated furcation rate, the analysis revealed that two non-coding mutations at the position of 28271 (g.a28271-/t) might be crucial for the high transmissibility of Alpha, Delta and Omicron VOCs. Both two mutations cause an A-to-T change at the core Kozak site of the *N* gene. The analysis also revealed that the non-coding mutations (g.a28271-/t) alone are unlikely to cause high viral transmissibility, indicating epistasis or multilocus interaction in viral transmissibility. A convergent evolutionary analysis revealed that g.a28271-/t, S:P681H/R and N:R203K/M occur independently in the three-VOC lineages, suggesting a potential interaction among these mutations. Therefore, this study unveils that non-synonymous and non-coding mutations could affect the transmissibility synergistically.

## Introduction

Three prevalent SARS-CoV-2 Variants of Concern (VOCs) were emerged in the last two years and draw the most attention for the epidemic waves that they caused (Fig. 1A). These VOCs were named as the Alpha, Delta and Omicron VOCs. The Alpha VOC, also known as SARS-CoV-2 lineage B.1.1.7, is a variant first detected in the UK ^1^ and has higher transmissibility than the preexisting variants in September 2020 ^2^. Its high transmissibility remains similar across different age, sex and socioeconomic strata ^3^. The Delta VOC was first identified in India in October 2020 ^4^ and has higher transmissibility ^5-7^. The Omicron VOC was first identified in Southern Africa in late November 2021 ^8^. Then the variant spreads more rapidly than the Delta variant ^9^ and has been detected in 76 countries globally in less than a month ^10^.

**Figure 1.**
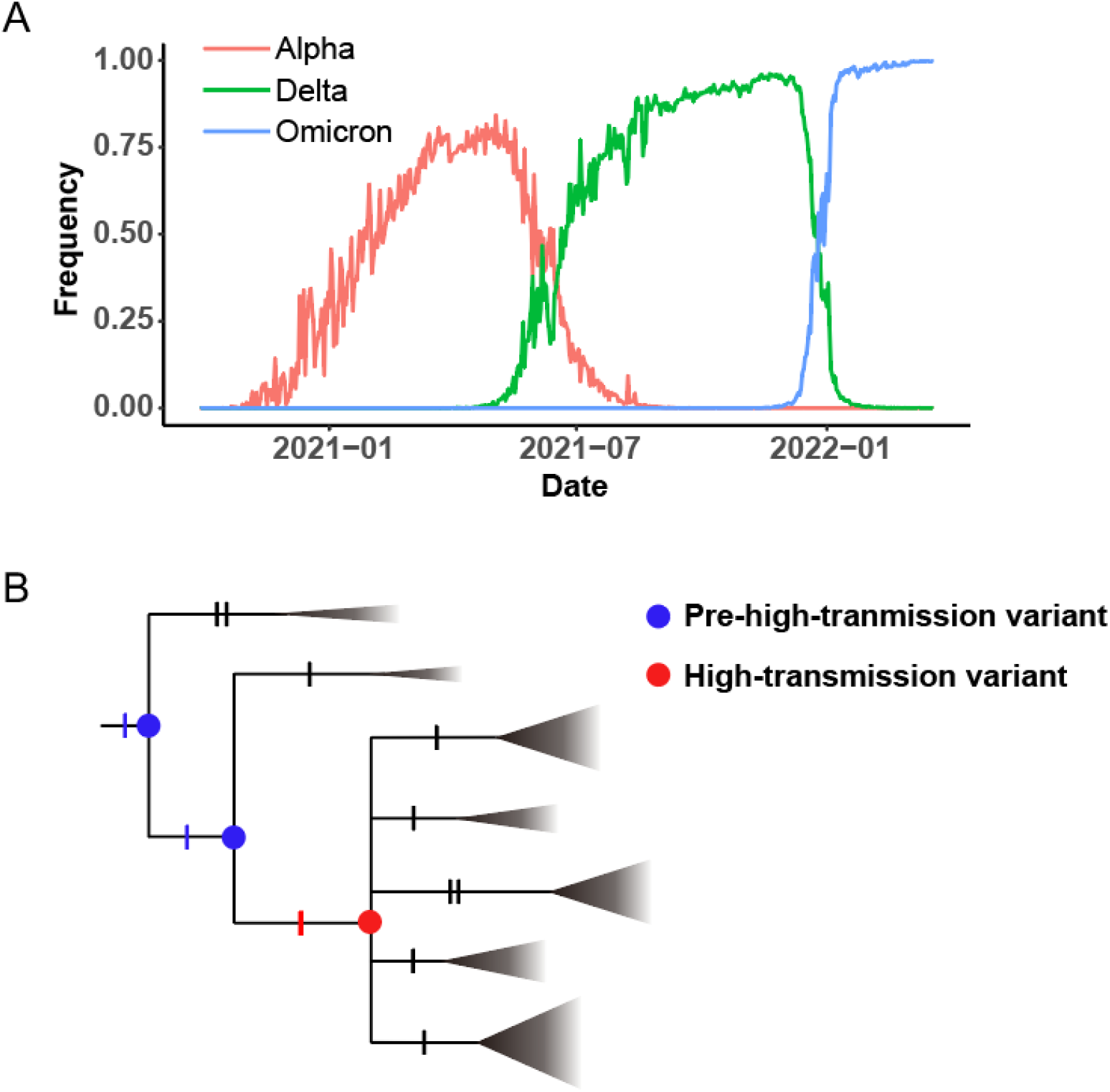
COVID-19 waves and a sudden appearance of high transmissibility model. (A) Three COVID-19 waves caused by the Alpha, Delta and Omicron VOCs. (B) Schematic diagram of a sudden appearance of high transmissibility model. The triangles represent collapsed clades and the size of the triangle represents the size of the clade. Each notch of branches represents a mutation. The last crucial mutation is indicated in red.

It is essential to uncover the key genetic changes that cause the high transmissibility of VOCs ^11,12^. All the three VOCs carry a large number of mutations. Comparing with the reference genomic sequence of SARS-CoV-2 (GenBank accession number: NC_045512.2) ^13^, the sequences of the Alpha, Delta, Omicron BA.1 and Omicron BA.2 variants have 21, 27, 52 and 51 amino-acid change mutations, respectively ^4^ while two, three, one and two non-coding mutations can be found ^14^. Moreover, different viral mutations are generally tightly linked since the virus lacks recombination occurred during the stage of meiosis in sexual species. Therefore, traditional population genetic methods, such as selective sweeps, may not reliably detect beneficial mutations ^15-19^ in SARS-CoV-2, and a novel approach is urgently needed.

In this study, we proposed a new pandemic-scale phylogenomic approach based on the sequential occurrence order of mutations and the instantaneously accelerated furcation rate to detect mutations crucial to transmissibility. Let us assume that the high transmissibility of a VOC is due to the emergence of a beneficial haplotype with multiple crucial mutations. Thus, the last (*i*.*e*., the most recent) crucial mutation is likely to increase the furcation rate in phylogenetic tree (Fig. 1B) because the VOC with a beneficial haplotype emerges. To identify the crucial coding and non-coding mutations for the three VOCs, we used the Coronavirus GenBrowser (CGB) ^14^ to analyze a tip-dated tree with 3,646,973 high-quality SARS-CoV-2 genomic sequences and the associated epidemiology metadata. The CGB is a free platform offering a panoramic vision of the transmission and evolution of SARS-CoV-2. The analysis revealed that two non-coding mutations at the position of 28,271 (g.a28271-/t) may be crucial for the high transmissibility of the three VOCs. Both two mutations cause an A-to-T change at the core Kozak site of the *N* gene. The analysis also revealed that the non-coding mutations (g.a28271-/t) alone are not associated with high viral transmissibility, indicating that the non-coding mutations may interact with other coding mutations to increase the viral transmissibility.

## Methods Data sources

The annotated evolutionary tree and evolutionary network data were obtained from the Coronavirus GenBrowser (CGB) ^14^ and VENAS ^20^. All sequence data of SARS-CoV-2 were obtained from the 2019nCoVR database ^21,22^, which is an integrated resource based on Global Initiative on Sharing All Influenza Data (GISAID) ^23,24^, National Center for Biotechnology Information (NCBI) GenBank ^25^, China National GeneBank DataBase (CNGBdb) ^26^, the Genome Warehouse (GWH) ^27^, and the National Microbiology Data Center (NMDC, https://nmdc.cn/). Two data versions of CGB (“data.2021-03-06” and “data.2022-04-14”) were used in this study, which contains 400,051 and 3,777,753 genomic sequences, respectively.

### Furcation rate to evaluate virus transmissibility

To evaluate virus transmissibility, we proposed to calculate furcation rate (FR) of nodes in a considered phylogenetic clade (Fig. 1B). Furcation rate of the *i*-th node (*FR*_*i*_) can be calculated using the following equation,

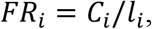

where *C*_*i*_ is the number of child nodes and *l*_*i*_ the branch length of the node with time in units. Since *FR*_*i*_ represents the furcation rate during the spanning time period of the branch, the furcation rate at time *t* (*FR*_*t*_) can be calculated by averaging the *FR*_*i*_ of branches that cover the time point in the considered phylogenetic clade. The estimated curve of *FR*_*t*_ can be smoothed to reduce the fluctuation. In this study, it was done by applying a sliding window with seven days in size.

### The effectiveness of mutations in improving viral transmissibility

To evaluate the effectiveness of a mutation in improving transmissibility, we tested whether two sibling clades with or without the mutation have similar viral transmissibility (Fig. 1B). The null hypothesis is that the two sibling clades have the same viral transmissibility. The genetic background of the two sibling clades has only two-mutation difference in the presented example (Fig. 1B). The alternative hypothesis is that the clade with the mutation (red) has higher viral transmissibility than the sibling clade without the mutation. The binomial probability was applied to test the null hypothesis and the test was one-tailed. Because the three VOCs occurred at different time points, the analysis for Alpha was based on the data version “data.2021-03-06” (*n* = 400,051) and the analysis of Delta and Omicron was based on the data version “data.2022-04-14” (*n* = 3,777,753), where *n* is the number of viral strains. The two datasets were obtained from the CGB ^14^.

### Calculation of *Rt*

Considering that the sequenced strains were sampled from the infected patients, the frequency of each variant at time *t* can be estimated from the CGB datasets and the number of new infections of a variant at time *t* (*I*_*V,t*_) can be approximated using the following equation,

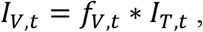

where *I*_*T,t*_ represents the total number of new infections at time *t* and *f*_*V,t*_ is the frequency of the variant at time *t*.

To reduce the fluctuation caused by the small frequency of the Alpha-like variant, we used a window size of 35 days to smooth its frequency curve before using it to calculate *Rt*. The total number of new infections (*I*_*T,t*_) in the United Kingdom was downloaded from the UK Coronavirus Dashboard (https://coronavirus.data.gov.uk/). Then the *Rt* was calculated for the Alpha and Alpha-like variants using the EpiEstim ^28^ with a mean (±SD) serial interval of 7.5± 3.4 days ^29^.

### Reappearance of g.a28271- and g.a28271t in the evolutionary tree

To examine the reappearance of mutations, we searched the non-coding mutations in the CGB evolutionary tree ^14^. We used the string “A28271-” to search g.a28271- and “A28271T” to search g.a28271t. To present the reappearance patterns of mutations, the data version “data.2021-04-14” (*n* = 3,777,753) of the CGB ^14^ was used. This data was also used to examine the frequency trajectory of g.a28271-/t when the Alpha, Delta and Omicron lineages were excluded.

### Identification of new canonical Alpha genomic sequence

The CGB was employed to identify a new canonical Alpha genomic sequence ^30^ that carries the deletion g.a28271- and all other Alpha characteristic mutations (Fig. 2A, Supplementary Table S1). We first selected all the strains in the Alpha (CGB84017.91425) clade that carries all the Alpha characteristic mutations including g.a28271-. Then we filtered the strains by date and only kept the strains collected before 1 November, 2020. Viral strains with extra mutations were ignored. Then the sequence with accession EPI_ISL_629703, as the suggested new canonical Alpha genomic sequence, is the first collected high-quality sequence without any extra mutations after the deletion g.a28271-occurred (Supplementary Fig. S1).

**Figure 2.**
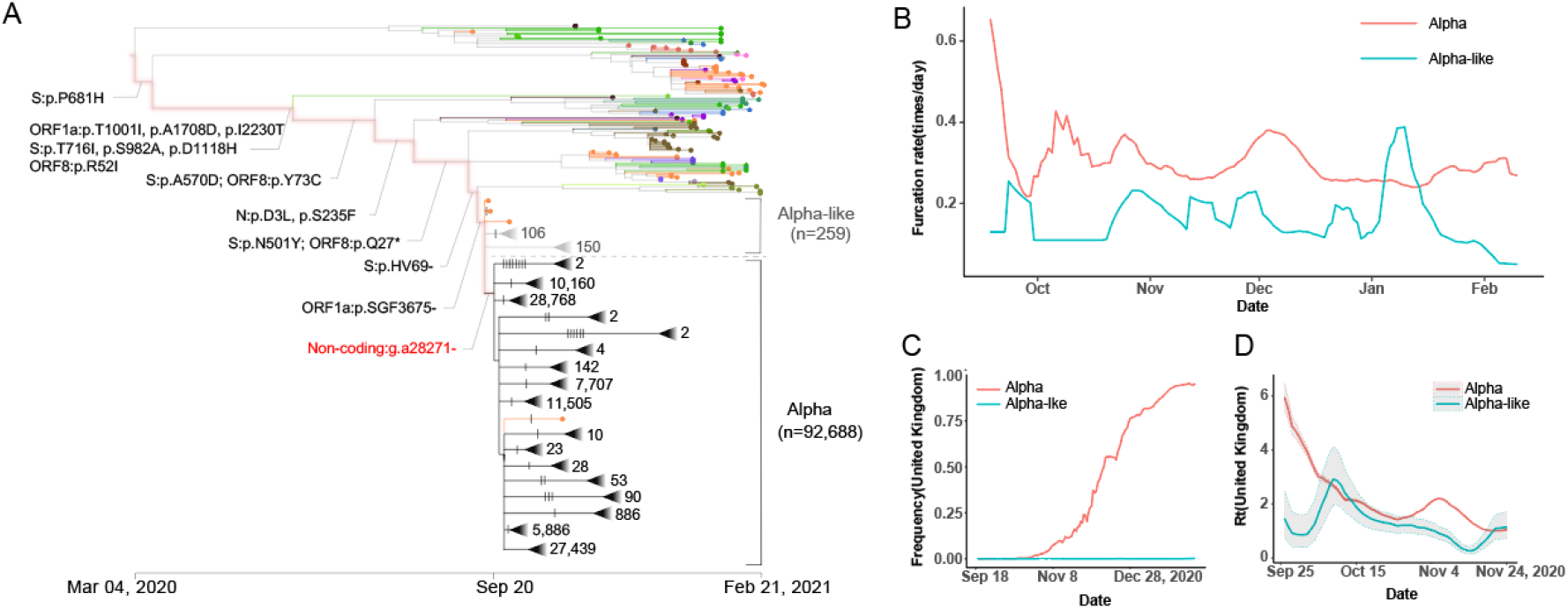
High transmissibility of Alpha compared with Alpha-like. (A) CGB evolutionary tree of SARS-CoV-2 lineage Alpha. The analysis was performed on 400,051 high-quality SARS-CoV-2 genomic sequences (data version “data.2021-03-06”) using the Coronavirus GenBrowser ^14^. The searchable CGB ID of the internal node with g.a28271-is CGB84017.91425, assigned by the CGB binary nomenclature system. The triangles represent collapsed clades. The size of collapsed clades was labeled after the collapse triangle. The mutations on the highlighted branches were labeled and the mutations on the Alpha and Alpha-like clades were marked by notches. The number of Alpha-like strains is 259, and the number of Alpha strains is 92,688. (B) The furcation rate of Alpha and Alpha-like. (C) The frequency trajectory of Alpha and Alpha-like in United Kingdom. (D) The Rt of Alpha and Alpha-like in United Kingdom. The grey area represents the 95% confidence interval.

## Results

### Instantaneously accelerated furcation rate and high transmissibility

It has been recognized that the phylogenetic tree is related to the transmission tree and different transmission patterns can be reflected in phylogenetic tree shapes ^31^. Thus furcation, as the process of lineage splitting ^32^, is associated with transmission. A variant with high transmissibility could cause more infections and has a greater chance to form new lineages in the examined evolutionary tree. These short and multifurcation internal nodes can be measured by the accelerated rate of furcation (see the details in Methods). Super-spreader transmission can also result in a multifurcation node, while it only affects the furcation rate of a single node. Thus, it can be distinguished from cases with high transmissibility.

Let us consider that a SARS-CoV-2 VOC has a beneficial haplotype with multiple crucial mutations. To explain the sudden appearance of its high transmissibility, we assume an instantaneous increase of fitness advantage as soon as the last crucial mutation occurs (Fig. 1B). In this model, we call variant with a subset of the beneficial mutations as pre-high-transmission variant, the furcation rate of which may remain nearly unchanged. However, as soon as the last crucial mutation occurs, the beneficial variant emerges and its transmissibility increases suddenly and causes an instantaneously accelerated furcation rate in phylogenetic tree.

### A crucial non-coding deletion in the Alpha VOC

The Alpha lineage was examined using the Coronavirus GenBrowser (CGB) ^14^. The sequential occurrence of the Alpha characteristic mutations was shown (Fig. 2A). To confirm the evolutionary path of the Alpha VOC, we also applied the VENAS ^20^ to obtain an evolution network of SARS-CoV-2 major haplotypes (Supplementary Fig. S2, Supplementary Table S1). The VENAS results are consistent with the CGB evolutionary tree.

The CGB evolutionary tree shows that many nodes remain to be bifurcating where the Alpha characteristic amino acid mutations occur (Fig. 2A). Multi-furcating nodes are frequently observed after a non-coding deletion (g.a28271-) occurs in the Alpha clade. The rate of furcation was calculated (Fig. 2B). The rate of furcation in the Alpha clade is higher than that in the Alpha-like clade. The Alpha-like clade indicates the viral strains lacking the non-coding deletion but carrying all other characteristic mutations of the Alpha VOC (Fig. 2A, Supplementary Table S1) ^30^, including all Alpha spike mutations ^33^. The Alpha clade carries all those characteristic mutations and the non-coding deletion. It was also found that the higher furcation rate of the Alpha clade lasted for five months, indicating that it is not due to super-spreader transmission.

The frequencies of Alpha and Alpha-like strains were calculated (Fig. 2C). It was found that, within six months, the global frequency of Alpha increases rapidly to over 80% while that of Alpha-like is no more than 1%. Moreover, the reproduction numbers (Rt) of Alpha and Alpha-like in the United Kingdom were estimated using the EpiEstim ^28^. The Rt of the Alpha lineage was generally larger than that of the Alpha-like lineage (Fig. 2D). The larger Rt estimates of the Alpha lineage indicates that the Alpha lineage has a transmission advantage over the Alpha-like. Therefore, the non-coding deletion g.a28271-may effectively increase the viral transmissibility.

The higher transmissibility of the Alpha lineage causes that the number of descendants in the Alpha lineage is highly significantly different with that in the Alpha-like lineage (*n* = 92,688 *vs* 259, *P*-value< 4.9 × 10^−324^). Since pooling data of viral sequences from different countries is likely to be biased due to complex differences in sampling with respect to either viral genome sequencing capacities or anti-contagion policies on the pandemic among the targeted countries ^34^. To address this problem, the numbers of descendants in the Alpha-like and Alpha lineages were pairwise compared for individual countries and continents, i.e., England (27 *vs* 76,871), Spain (30 *vs* 712), Switzerland (8 *vs* 1,332), Germany (2 *vs* 570), USA (8 *vs* 1,028), Australia (1 *vs* 58), South America (1 *vs* 22), Africa (1 *vs* 86), and Asia (3 *vs* 642). The transmissibility of strains without or with the non-coding deletion is significantly unequal (Supplementary Table S2, *P*-value≤ 2.74 × 10^−6^). The same highly significant statistics was observed in 10 more countries, such as India and Italy. Moreover, the same conclusion holds when considering different gender and age groups (Supplementary Tables S3, S4). Therefore, the significantly difference in transmissibility between the Alpha and Alpha-like clades has been observed which is likely due to the non-coding deletion g.a28271-.

### The crucial non-coding deletion in the Delta VOC

The non-coding deletion (g.a28271-) was also found in the Delta VOC (Fig. 3A). We used a recent data version (data.2022-04-14) to analyze this VOC since it became prevalent globally after the Alpha VOC. It was observed that the number of furcation increases as soon as the non-coding deletion (g.a28271-) occurs in the Delta lineage (Fig. 3A), similar with the case in the Alpha lineage. The rate of furcation in the Delta lineage is generally higher than that in the Delta-like lineage (Fig. 3C). The Delta-like lineage represents the viral strains lacking the non-coding deletion but carrying all other characteristic mutations of the Delta VOC (Fig. 3A).

**Figure 3.**
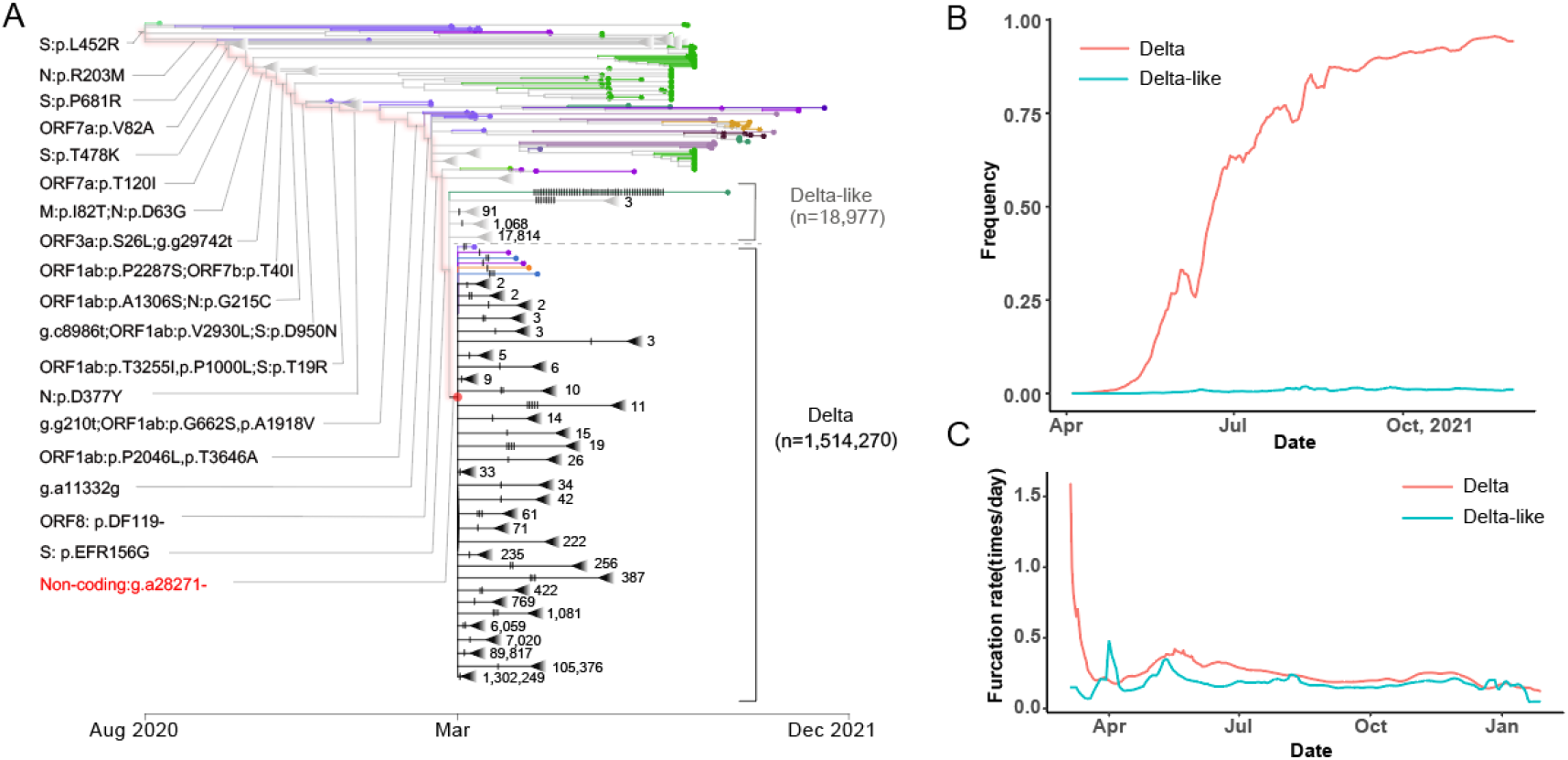
High transmissibility of Delta compared with Delta-like. (A) CGB evolutionary tree of SARS-CoV-2 lineage Delta. The analysis was performed on 3,777,753 high-quality SARS-CoV-2 genomic sequences (data version “data.2022-04-14”) using the Coronavirus GenBrowser ^14^. The searchable CGB ID of the internal node with g.a28271-is CGB531065.736525, assigned by the CGB binary nomenclature system. The triangles represent collapsed clades. The size of collapsed clades was labeled after the collapse triangle. The mutations on the highlighted branches were labeled and the mutations on the Delta and Delta-like clades were marked by notches. The number of Delta-like strains is 18,977, and the number of Delta strains is 1,514,270. (B) The frequency trajectory of Delta and Delta-like. (C) The furcation rate of Delta and Delta-like.

The global frequencies of Delta and Delta-like strains were calculated (Fig. 3B). It reveals that the global frequency of Delta increases rapidly to over 90% within six months while that of Delta-like remains to be around 1%. The higher transmissibility of the Delta lineage also causes that the number of its descendants is highly significantly different with that in the Delta-like lineage (*n* = 1,514,270 *vs* 18,977, *P*-value< 4.9 × 10^−324^). All these results support the modulation of non-coding deletion g.a28271-on viral transmissibility.

### A crucial non-coding mutation in the Omicron VOC

The Omicron VOC first reported in South Africa in November 2021 ^8^ that evolved into multiple sub-lineages, such as BA.1 and BA.2. The CGB evolutionary tree shows that the most Omicron samples also carry a mutation at position 28271 (g.a28271t) (Fig. 4A). The mutation occurred independently in the two sub-lineages (BA.1 and BA.2). Therefore, to examine the effects of the mutation on the viral transmissibility, the furcation rate was calculated for the lineages with and without the mutation. To keep the consistence, the term of Omicron stands for strains with the mutation g.a28271t while Omicron-like indicates strains without the mutation in the considered lineage (Fig. 4A). It was found that the Omicron lineage appears to have a higher furcation rate than the Omicron-like lineage, indicating that the mutation g.a28271t plays a key role in the viral transmissibility. Different from the situation in Alpha and Delta, a delayed increase of furcation rate was observed (Fig. 4), which suggests that g.a28271t may interact with other mutations to form a beneficial haplotype.

**Figure 4.**
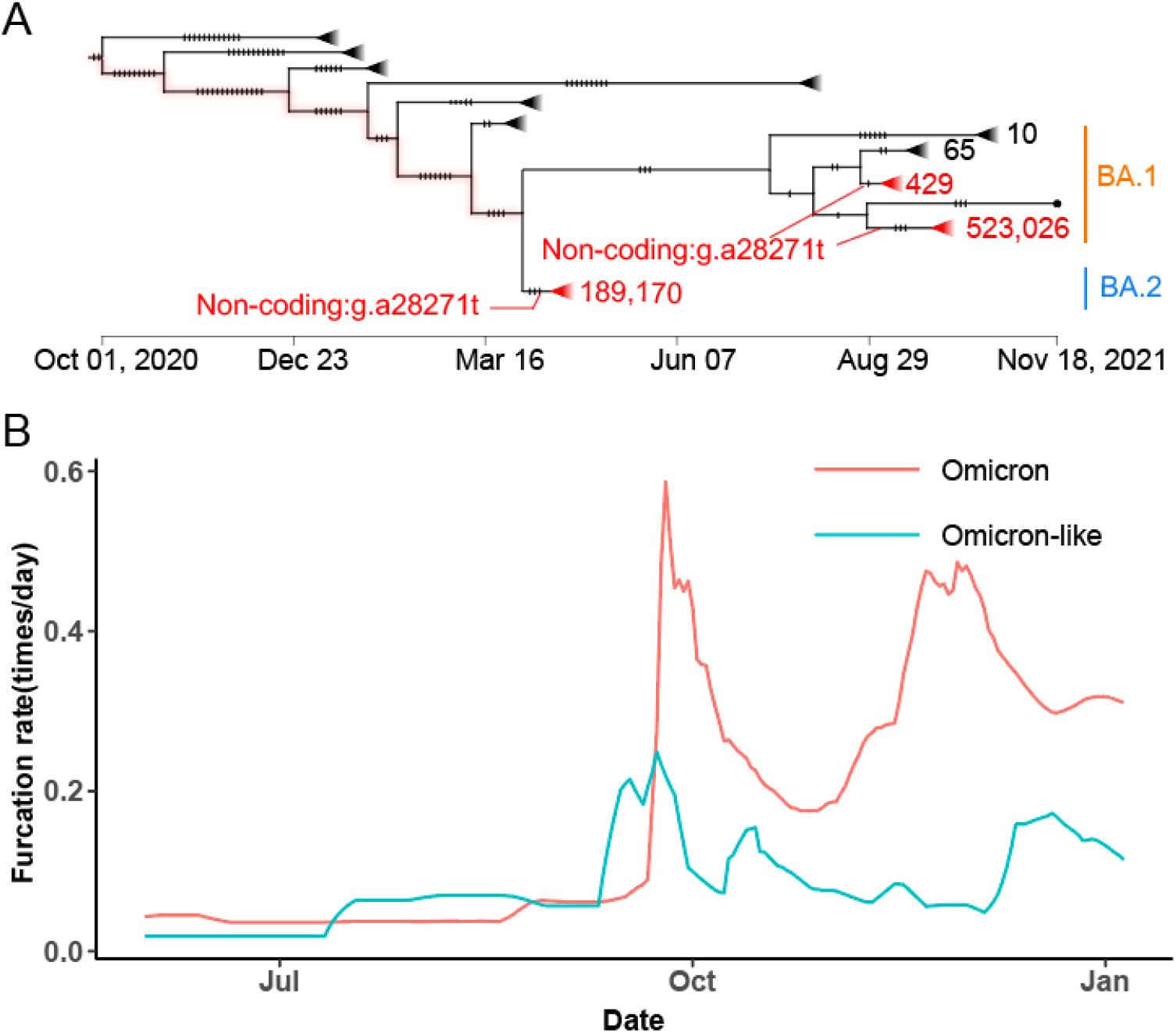
High transmissibility of Omicron compared with Omicron-like. (A) CGB evolutionary tree of SARS-CoV-2 lineage Omicron. The analysis was performed on 3,777,753 high-quality SARS-CoV-2 genomic sequences (data version “data.2022-04-14”) using the Coronavirus GenBrowser ^14^. The CGB ID of the internal nodes with g.a28271t are CGB3145480.3251694 and CGB2437061.2958631 for BA.1, CGB3053091.3265861 for BA.2. The triangles represent collapsed clades. The size of collapsed clades was labeled after the collapse triangle. The mutations were marked by notches. (B) The furcation rate of Omicron and Omicron-like.

### The non-coding mutations changes the core Kozak site of *N* gene

The base 28,271 is located at the third base upstream of the start codon of the *N* gene. It is the core Kozak site of the gene (Supplementary Fig. S3). The Kozak sequence is a short sequence around the start codon and functions as the protein translation initiation site in higher eukaryotes ^35^. Nucleotides in each site of Kozak sequence influence the translational efficiency, especially the positions -3 and +4. The translational efficiency of the gene will be higher with A or G in position -3 and G in position +4 ^35^. The position 28,271 is the core position -3 of Kozak sequence of the *N* gene. The g.a28271-deletion makes t28,270 to slip one base and changes the Kozak context of gene *N* from a suboptimal one (A at -3, T at +4) to an undesirable one (T at -3, T at +4) (Supplementary Fig. S3) ^35^. It was expected that the mutation g.a28271t may produce a similar effect by making the Kozak context of gene *N* change to an undesirable one (T at -3, T at +4). When the homological site of the SARS-CoV genome was mutated to another undesirable one (C at -3, T at +4), the expression of N protein was reduced and the translation of ORF9b protein increased ^36^. The ORF9b protein was encoded by an alternative reading frame within the ORF of gene *N* and the protein was found to be translated via a leaky ribosomal scanning mechanism ^36^. It has been found that SARS-CoV-2 ORF9b has an interferon (IFN) antagonistic activity and suppresses type I interferon (IFN-I) responses ^37,38^. Therefore, the non-coding mutations may strengthen the interference of ORF9b on IFN-1 responses by changing the core Kozak site of *N* gene.

### The non-coding mutations alone do not increase viral transmissibility

The non-coding mutations g.a28271- and g.a28271t were found to be crucial for the high transmissibility of SARS-CoV-2. It was investigated whether these two mutations function alone or co-act with other crucial mutations by considering recurrent g.a28271- and g.a28271t mutations (including mutations due to recombination). The non-coding mutations have appeared multiple times in the evolution of SARS-CoV-2 (Supplementary Fig. S4), the frequency trajectories of the two mutations were checked when the Alpha, Delta and Omicron lineages were excluded. The first sample that carries one of the mutations appeared on June 16, 2020 (EPI_ISL_732537, with g.a28271-) and the frequency of the two mutations generally remains much less than 10% in the next six months (Supplementary Fig. S5). Therefore, the two non-coding mutations require epistasis or multilocus interaction with other mutations to increase the viral transmissibility, consistent with the observations in the Omicron VOC.

### The convergent evolution of the Alpha, Delta and Omicron VOCs

To identify mutations that interact with g.a28271-/t, the convergent evolution of the Alpha, Delta and Omicron VOCs may provide important clues. The three VOCs diverged in January, 2020, and evolved independently in the background of the D614G substitution (Fig. 5A). Thus the mutations of the three VOCs were screened in the phylogenetic tree. The two noncoding mutations (g.a28271- and g.a28271t) evolved convergently in the three VOCs (Fig. 5B). Moreover, another two non-synonymous mutations (S:P681H/R and N:R203K/M) were found to occur independently in the lineages.

**Figure 5.**
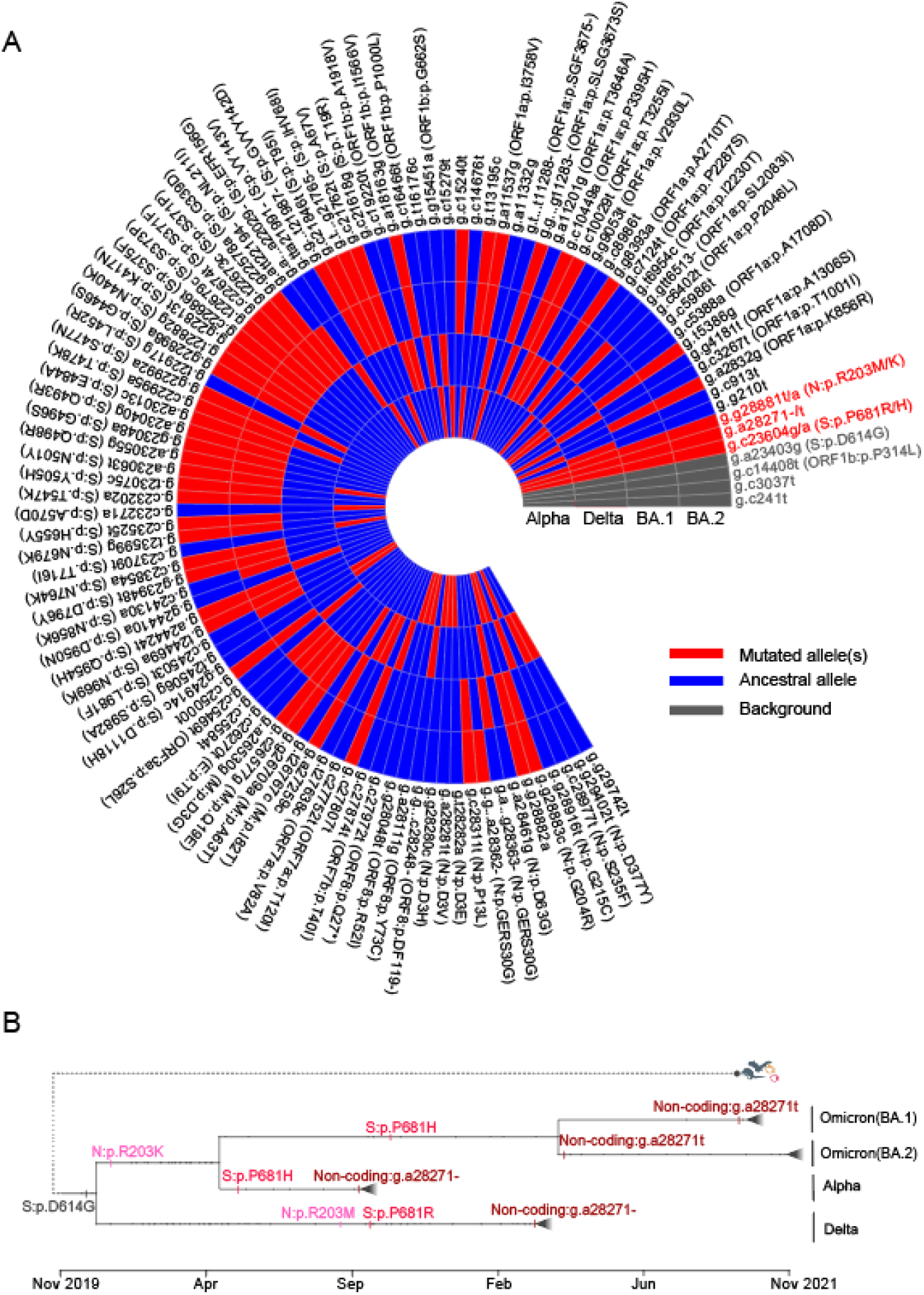
The convergent evolution of the Alpha, Delta and Omicron VOCs. (A) Heatmap showing the presence of mutations in Alpha, Delta and Omicron (BA.1, BA.2) VOCs. In the heatmap, four background mutations are in grey, mutated alleles are in red and ancestral alleles in blue. (B) Schematic diagram of convergent evolution of Alpha, Delta and Omicron VOCs. Mutations under convergent evolution are labeled, the occurring time of mutations are inferred by the CGB ^14^.

Position 681 in the S protein is immediately upstream of the furin cleavage site ^39^. The mutation S:P681H occurred independently in the Alpha and Omicron VOCs (Fig. 5B). It has been found that the mutated spike (681H) can be partially cleaved when a protease inhibitor is added, while the un-mutated spike (P681) is nearly uncleaved. This result suggests that the P681H mutation may increase furin cleavage ^40^. The mutation S:P681R, occurred in Sep, 2020 in the Delta VOC, also has the similar effects ^41^. Therefore, both the mutations S:P681H/R may be important for the furin cleavage and viral entry.

The N protein is required for replication and packaging. Position 203 is in the linker region of the N protein. The mutation N:R203K occurred on the common ancestor of Alpha and Omicron VOCs in Feb, 2020 (Fig. 5B). Its adjacent mutation N:G204R also occurs on the two VOCs (Fig. 5A). It has been found that the double-mutated strain (203K/204R) has a replication advantage over the wild-type (R203/G204) ^42^. Similarly, the mutation N:R203M occurred in Sep, 2020, before the Delta VOC emerged (Fig. 5B). The mutation N:R203M can enhance replication in lung epithelial cells ^43^. Therefore, both the mutations N:R203K/M may be crucial for replication.

## DISCUSSION

It is generally difficult to map causal mutations with fitness advantage along the SARS-CoV-2 genome using traditional population genetic methods, such as selective sweep ^15,18,19^, because of the extreme rarity of recombination in asexual population. All mutations will be tightly linked and the methods to detect selective sweep actually are not developed to be applied to viral genomes. For the same reason, machine-learning based approaches ^16^ may also face the same challenge. Therefore, we analyze the sequential occurrence order of mutations and the change of furcation rate in the pandemic-scale phylogenetic tree of SARS-CoV-2 to map viral mutations with fitness advantage. We find that the non-coding g.a28271-mutation may play a crucial role in the high transmissibility of the Alpha and Delta VOCs, and g.a28271t for the Omicron VOC. Both the mutations cause the A-to-T change at the core Kozak site of the gene *N*. The Alpha-like and the Alpha strains encode the same spike protein; however, these strains appear to have different viral transmissibility. The Alpha and the Alpha-like spikes improve the angiotensin-converting enzyme 2 (ACE2) affinity for about 5-fold, comparing with the D614G spike ^33^. However, our analysis shows that the increase of ACE2 affinity alone may not cause the high transmissibility of the Alpha VOC when the non-coding deletion g.a28271-is lack.

Sequence with accession EPI_ISL_601443 was previously recommended to be the canonical Alpha genomic sequence ^30^. However, it does not carry the crucial non-coding deletion g.a28271-. Therefore, to investigate the viral transmissibility, we suggest using the sequence with accession EPI_ISL_629703 as the canonical Alpha genomic sequence (collected 21 October, 2020, in the UK) (Supplementary Fig. S1).

Genomic mutations related to the transmissibility of a pandemic etiological pathogen such as SARS-CoV-2 is complex and difficult to be revealed merely *via* genetic analysis with limited and incomplete supporting data of epidemiology.

However, this study unveils that non-synonymous and non-coding mutations could affect the transmissibility synergistically as a beneficial haplotype. Overall, our analyses indicate that the non-coding A-to-T kozak site change of the gene *N* could be crucial for the high transmissibility of SARS-CoV-2 and may interact with mutations on the spike S1/S2 cleavage site and mutations that influence virus replication.

## Acknowledgments

We thank Wolfgang Stephan for discussing on the issue of selective sweep and the researchers who generated and deposited sequence data of SARS-CoV-2 in GISAID, GenBank, CNGBdb, GWH, and NMDC. This work was supported by grants from the Strategic Priority Research Program of the Chinese Academy of Sciences (Grant No. XDB38030100), the National Key Research and Development Project (Grant Nos. 2020YFC0847000, 2021YFC0863300, and 2020YFC0845900), the National Natural Science Foundation of China (Grant No. 91531306), and the Shandong Academician Workstation Program #170401 (to G.P.Z.).

## Data Availability

All the SARS-CoV-2 data can be obtained from the Coronavirus GenBrowser and VENAS. Raw data were deposited on Mendeley at https://data.mendeley.com/datasets/tbbxjy3gyr/draft?a=1be355b0-e347-4599-89dd-19c950051f02. Other data are available from the corresponding authors upon request.

## Authors’ contributions

Conceptualization, J.Y., G.Z., D.Y., Y.H.P., B.S., Y.Z., G.P.Z., Y.L., H.L.; Data Analysis, J.Y., G.Z., D.Y., R.C., X.W., Y.L., C.Y., X.S.; Writing, J.Y., G.Z., D.Y., R.C., Y.L., Y.H.P., C.Y., X.S., Y.Z., G.P.Z., H.L.; Supervision & Funding Acquisition, G.Z., Y.Z., G.P.Z., Y.L., H.L.

