## Supplementary Appendix for "A non-coding A-to-T Kozak site change related to the transmissibility of Alpha, Delta and Omicron VOCs"

[†]These authors contributed equally.

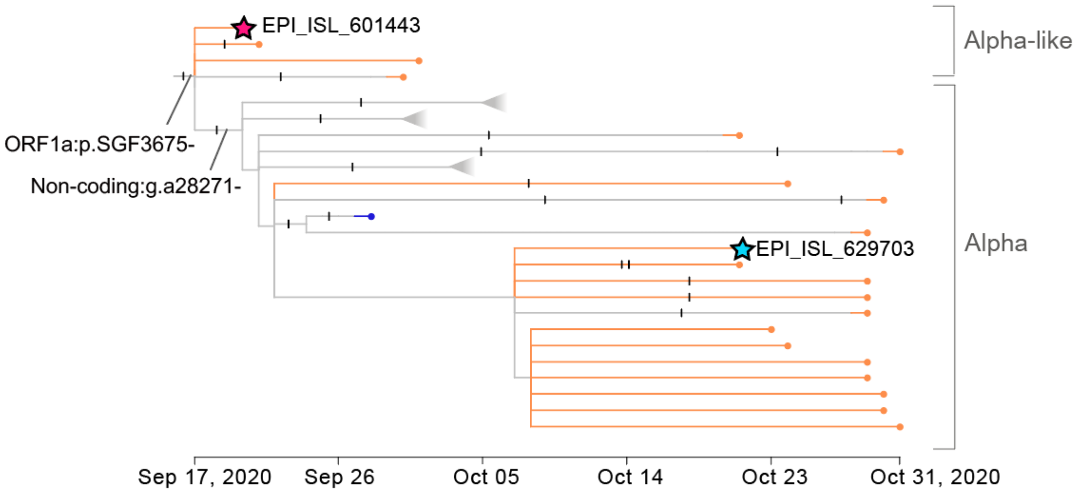


**Figure S1. Suggested canonical Alpha genomic sequence.**

The current canonical Alpha genomic sequence was marked in red star. The new canonical Alpha genomic sequence we suggested was in blue star. Each notch of the branches represents a mutation. The branches with no mutations were highlighted. The blue-star sequence is the first collected high-quality sequence without any extra mutations after the deletion g.a28271- occurred.

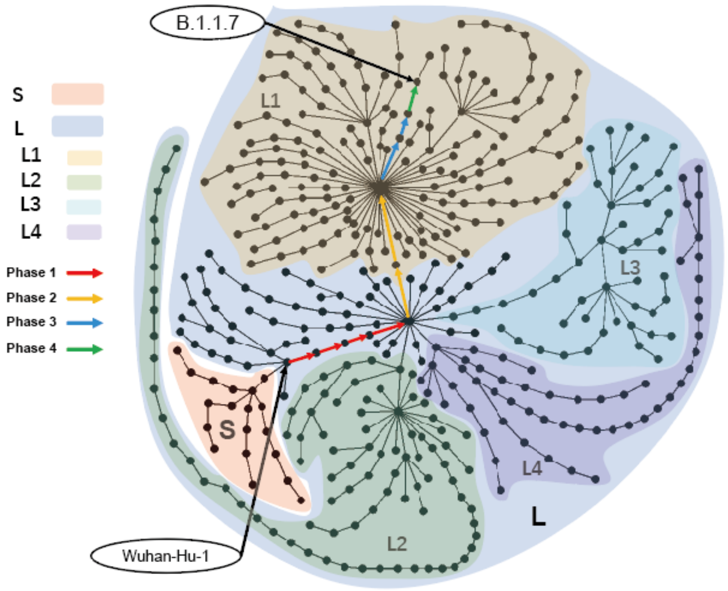


**Figure S2. VENAS evolution network of SARS-CoV-2 by January 14, 2021.**

The dots represent the major genome types of SARS-CoV-2, and the lines between the dots are the evolutionary path formed by the combination of variants; the color shades represent the clades and subclades formed by genome types, where the L1 subclade is shaded in yellow; the L2 in green; the L3 in cyan, and the L4 in purple. The L/S naming system follows the previous study [1]. The color arrows mark the evolutionary path from the most recent common ancestor of SARS-CoV2 to the Alpha linage, and four phases are indicated in different colors.

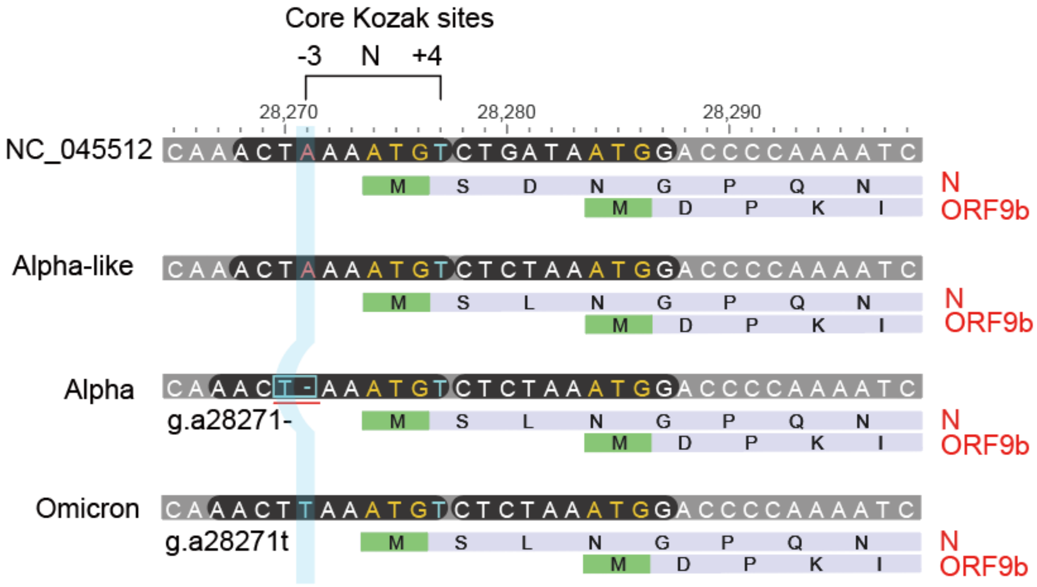


**Figure S3. A non-coding A-to-T Kozak site change related to the transmissibility of VOCs.**

Mutations in position 28271 change the core Kozak sites of N and ORF9b genes. The two positions -3 and +4 have the dominant influence [2]. The grey bars are the nucleotide sequences of the variants. Two functional genes are presented under each sequence. Start codons are shown in green. The N and ORF9b genes with their amino acid sequences are colored in light purple. Sites that mutations happened are covered in light blue rectangle. The optimal Kozak sites are colored in red and non-optimal ones in light blue.

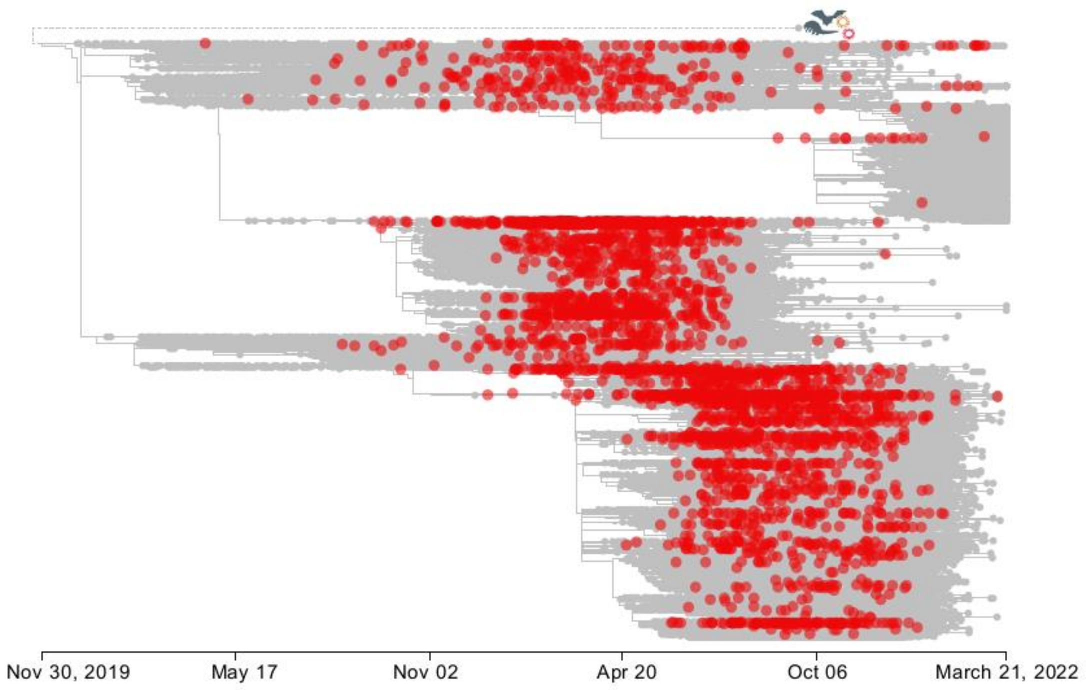


**Figure S4. Reappearance of g.a28271- or g.a28271t in an evolutionary tree with 3,777,753 high-quality SARS-CoV-2 genomic sequences.**

The red dots indicate branches with g.a28271- or g.a28271t, it can be recurrent mutations or mutations due to recombination.

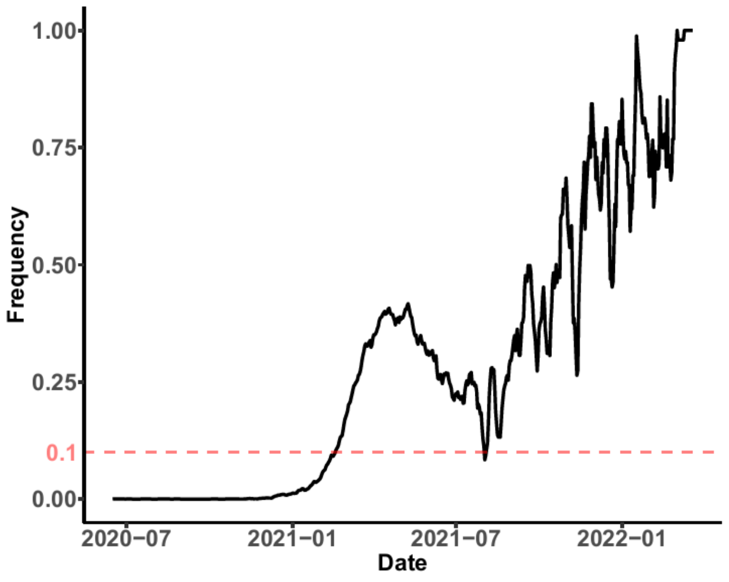


**Figure S5. Frequency trajectory of g.a28271- and g.a28271t when the Alpha, Delta and Omicron lineages were excluded.**

Samples with g.a28271- or g.a28271t are counted when the Alpha, Delta and Omicron lineages were excluded.

**Table S1. Mutations of Alpha lineage.**

| Evolutionary phase | Nucleotide mutation | Amino-acid mutation | Type |
|---|---|---|---|
| | g.c3267t | ORF1a:p.T1001I | SNV |
| | g.c5388a | ORF1a:p.A1708D | SNV |
| | g.t6954c | ORF1a:p.I2230T | SNV |
| | g. tctggtttt11288- | ORF1a:p.SGF3675- | INDEL |
| | g.tacatg21765- | S:p.HV69- | INDEL |
| | g.a23063t | S:p.N501Y | SNV |
| Phase4 | g.c23271a | S:p.A570D | SNV |
| | g.c23604a | S:p.P681H | SNV |
| (Characteristic | g.c23709t | S:p.T716I | SNV |
| mutations of | g.t24506g | S:p.S982A | SNV |
| Alpha) | g.g24914c | S:p.D1118H | SNV |
| | g.c27972t | ORF8:p.Q27* | SNV |
| | g.g28048t | ORF8:p.R52I | SNV |
| | g.a28111g | ORF8:p.Y73C | SNV |
| | g.a28271- † | Non-coding | INDEL |
| | g.gat28280cta | N:p.D3L | SNV |
| | g.c28977t | N:p.S235F | SNV |
| Phase3 | g.tta21991- | S:p.Y144- | INDEL |

| | | | |
|---|---|---|---|
| Phase2 | g.ggg28881aac | N:p.RG203KR | SNV |
| Phase1 | g.a23403g | S:p.D614G | SNV |
| | g.c14408t | ORF1ab:p.P4715L | SNV |

*Note*: The non-coding deletion is recommended to be one of characteristic mutations in this study. The deletion is not carried by the canonical Alpha genomic sequence (EPI_ISL_601443) [3].

**Table S2. The number of Alpha-like and Alpha strains in different countries and continents.**

| Country/continent* | The number of strains | | *P*-value |
|---|---|---|---|
| | Alpha-like | Alpha | |
| England | 27 | 76,871 | $< 4.9 \times 10^{-324}$ |
| Spain | 30 | 712 | $1.16 \times 10^{-170}$ |
| Switzerland | 8 | 1,332 | $< 4.9 \times 10^{-324}$ |
| Germany | 2 | 570 | $1.06 \times 10^{-167}$ |
| USA | 8 | 1,028 | $4.35 \times 10^{-293}$ |
| Australia | 1 | 58 | $1.02 \times 10^{-16}$ |
| Norway | 1 | 210 | $6.41 \times 10^{-62}$ |
| Denmark | 1 | 4,494 | $< 4.9 \times 10^{-324}$ |
| India | 2 | 16 | $5.84 \times 10^{-4}$ |
| Ireland | 2 | 897 | $9.55 \times 10^{-266}$ |
| France | 2 | 1,059 | $2.27 \times 10^{-314}$ |
| Sweden | 16 | 182 | $3.56 \times 10^{-37}$ |
| Finland | 26 | 198 | $2.60 \times 10^{-34}$ |
| Austria | 28 | 242 | $4.85 \times 10^{-44}$ |
| Italy | 29 | 734 | $5.33 \times 10^{-178}$ |
| Belgium | 72 | 1,230 | $4.52 \times 10^{-273}$ |
| South America | 1 | 22 | $2.74 \times 10^{-6}$ |
| Africa | 1 | 86 | $5.62 \times 10^{-25}$ |

| | | | |
|---|---|---|---|
| Asia | 3 | 642 | $3.05 \times 10^{-187}$ |

*Countries/continents with more than 10 viral strains (Alpha-like and Alpha).

**Table S3. The number of Alpha-like and Alpha strains in different gender groups.**

| Gender | The number of strains | | *P*-value |
|---|---|---|---|
| | Alpha-like | Alpha | |
| Male | 102 | 2,960 | $< 4.9 \times 10^{-324}$ |
| Female | 92 | 3,145 | $< 4.9 \times 10^{-324}$ |
| Unknown | 65 | 86,583 | $< 4.9 \times 10^{-324}$ |

**Table S4. The number of Alpha-like and Alpha strains in different age groups.**

| Age group | The number of strains | | *P*-value |
|---|---|---|---|
| | Alpha-like | Alpha | |
| 0-10 | 13 | 357 | $1.31 \times 10^{-88}$ |
| 11-20 | 19 | 568 | $4.86 \times 10^{-142}$ |
| 21-30 | 40 | 935 | $6.20 \times 10^{-223}$ |
| 31-40 | 34 | 968 | $4.80 \times 10^{-239}$ |
| 41-50 | 31 | 889 | $6.21 \times 10^{-220}$ |
| 51-60 | 28 | 879 | $1.29 \times 10^{-220}$ |
| 61-70 | 15 | 532 | $1.61 \times 10^{-136}$ |
| 71-80 | 8 | 329 | $1.36 \times 10^{-86}$ |
| 81-90 | 3 | 352 | $1.01 \times 10^{-100}$ |
| 91-100 | 1 | 114 | $2.77 \times 10^{-33}$ |
| Unknown | 67 | 86,765 | $< 4.9 \times 10^{-324}$ |
